# KusakiDB v1.0: a novel approach for validation and completeness of protein orthologous groups

**DOI:** 10.1101/2020.11.09.373753

**Authors:** Andrea Ghelfi, Yasukazu Nakamura, Sachiko Isobe

## Abstract

Plants have quite a low coverage in the major protein databases despite their roughly 350,000 species. Moreover, the agricultural sector is one of the main categories in bioeconomy. In order to manipulate and/or engineer plant-based products, it is important to understand the essential fabric of an organism, its proteins. Therefore, we created KusakiDB, which is a database of orthologous proteins, in plants, that correlates three major databases, OrthoDB, UniProt and RefSeq. KusakiDB has an orthologs assessment and management tools in order to compare orthologous groups, which can provide insights not only under an evolutionary point of view but also evaluate structural gene prediction quality and completeness among plant species. KusakiDB could be a new approach to reduce error propagation of functional annotation in plant species. Additionally, this method could, potentially, bring to light some orthologs unique to a few species or families that could have evolved at a high evolutionary rate or could have been a result of a horizontal gene transfer.

**Availability and Implementation:** The software is implemented in R. It is available at http://pgdbjsnp.kazusa.or.jp/app/kusakidb and at https://hub.docker.com/r/ghelfi/kusakidb under the MIT license.

**Supplementary information:** Supplementary data are available at *Bioinformatics* online.

## 1. Introduction

Plants have quite a low coverage in the major protein databases despite their roughly 350,000 species (Vellend, et al., 2017), and environmental and economic impacts, such as discussed in the Paris Agreement (UNFCCC, 2015) and the Global Bioeconomy Summit (GBS, 2015; GBS, 2018). Moreover, the agricultural sector is one of the main categories in bioeconomy (FAO, 2019), in 2013, biology-based industries accounted for 17 million jobs and more than US$ 2.2 trillion annually in the European Union (El-Chichakli, 2016).

In order to manipulate and/or engineer plant-based products, it is important to understand the essential fabric of an organism, its proteins. The three major protein databases, UniProt (Consortium, 2018), OrthoDB (Kriventseva, et al., 2019) and RefSeq (O’Leary, et al., 2016), have drawn our attention by the amount and type of information they maintain. UniProt arrange annotation layers such as GO (The Gene Ontology Consortium, 2019), EC (Bairoch, 2000), PFAM (Finn, et al., 2014), InterPro (Mitchell, et al., 2019) and KEGG (Kanehisa, et al., 2017). OrthoDB provides annotation of orthologous proteins, and RefSeq has the largest database of proteins available nowadays.

Yet some questions still may be raised regarding plant species (Viridiplantae). First, how could we evaluate the conservation of orthologous groups (OGs) within, for example, family level in OrthoDB? Second, could we increase the number of species in each OG by correlating all protein sequences in UniProt with OrthoDB? Third, which orthologs from OrthoDB are missing in UniProt that are found in RefSeq? Fourth, being aware that mis-annotation of molecular function in public databases continues to be a significant problem (Schnoes, et al., 2009) is it possible to assert the existence of an OG?

To address these questions, we created KusakiDB, which is a database of orthologs proteins, in plants, that correlates three major databases, OrthoDB, UniProt and RefSeq. It provides a validation tag for OGs based on a physical existence of at least one protein (or transcript) in each OG. Furthermore, KusakiDB has an OG assessment and OG management tools to compare OGs, within its own data but also user’s data, which can provide insights not only under an evolutionary point of view but also evaluate structural gene prediction quality and completeness among plant species. Comparisons of OGs among species within a family may disclose potential divergent annotation, which could be a new approach to reduce error propagation of functional annotation in plant species. Additionally, this method could, potentially, bring to light some orthologs unique to a few species or families that could have evolved at a high evolutionary rate or could have been a result of a horizontal gene transfer.

## 2. Material and Methods

### Cluster enrichment

KusakiDB performs a novel approach to enrich OGs by aligning protein sequence in OrthoDB-Viridiplantae sequences to protein sequences in UniProt-Viridiplantae and RefSeq-Viridiplantae, using sequence identity of 50 % and 80 %, respectively (Figure 1-A). Therefore, KusakiDB established a correlation between UniProt and RefSeq entries via OrthoDB cluster groups and resulted increase the number of plant species relating to the OGs from 117 (OrthoDB) to 126,030 in 814 families and 2 Phyla (Streptophyta and Chlorophyta). UniProt entries are investigated in prior to RefSeq entries. Fragmented sequences with flag “Fragmented” in UniProt were removed. RefSeq were used primary for KusakiDB validation, in 14,787 sequences that aligned with OrthoDB without any correspondent from UniProt. An HMMER profile (Eddy, 2011) version v3.3.1 (http://hmmer.org/) was built for each OG in OrthoDB, these profiles were used to validate the enriched clusters in KusakiDB with hmmsearch (HMMER v3.3.1), only entries that had E-value, global and first best domain, lower than 10^−3^ were considered. KusakiDB sequences were clustered, using UCLUST algorithm (Edgar, 2010) with USEARCH v11.0.667 (i86linux64), to 99% to remove redundancy.

**Figure 1.**
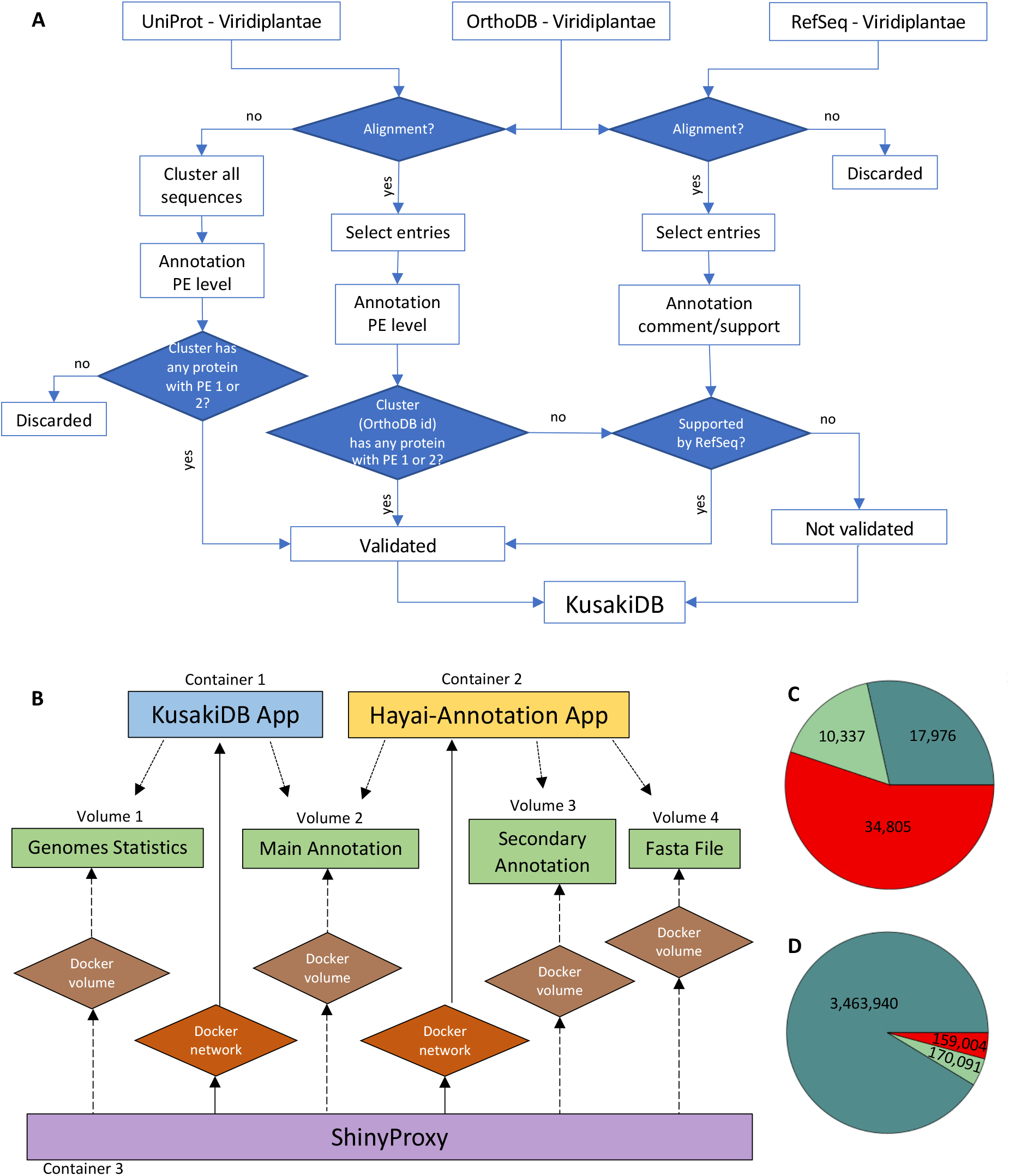
**A**: KusakiDB validation flowchart. Sequences from UniProt were aligned with OrthoDB using 50% of sequence identity. Sequences from RefSeq were aligned with OrthoDB with 80% of sequence identity. In both alignments were considered query length and target length 75%. Sequences from UniProt that did not align with OrthoDB were clustered with UCLUST using the criteria: sequence identity 50%, and query and target length 50%. Validation were provided by Protein Existence (PE) level in UniProt and ‘comment’ or ‘support’ in RefSeq; **B**: Overview of KusakiDB and Hayai-Annotation docker applications with ShinyProxy. **C:**KusakiDB unique OGs per validation status. **D:**KusakiDB all entries per validation status. Legends for **C**and **D**: (dark green: validated with UniProt, light green: validated with RefSeq, red: not validated).

### KusakiDB validation

KusakiDB validates an OG based on the physical existence of a protein, or transcript, sequence in each OG. The OG cluster is considered “validated” if there is at least one protein in the cluster with protein existence (PE) level 1 or 2 (existence at a protein level or transcript level, respectively) on UniProt entries or has any type of “support” of protein existence (mRNA or EST) or has either the tag “Reviewed” or “Validated” on RefSeq entries.

### OG Assessment Analysis

OG assessment calculates the total number of OGs per species, total number of species per family, percentage of validated entries “KusakiDB validated”, and median of OGs within a family “Median Family” and within all species “Median Species”. Same calculations can be performed with user’s data. For further information see GitHub.

Median of OGs within a family is the median of the frequency of unique OG per species within each family, in percentage. Median of OGs within all species is the median of the frequency of unique OG per species within all species, in percentage (Suppl. File 1).

### OG Management and OG Management User Data Analysis

OG Management tool shows the frequency of OGs per species and per protein name that fits in the parameters selected by users. The parameters are KusakiDB validation, total number of species in a family, presence of unique OG per species within a family (in percentage) and presence of unique OG per species within all species (in percentage). Same filter can be performed with user’s data. For further information please see GitHub.

### KusakiDB App Design

KusakiDB App is an R-package deployed using ShinyProxy in a docker container. The whole system contains four docker volumes and three containers (Figure 1-B).

## 3. Results and Discussion

A total of 63,118 unique OGs were registered in KusakiDB. The distribution of unique OGs in KusakiDB showed 28.5% were validated by UniProt, 16.4% were validated by RefSeq and 55.1% were not validated (Figure 1-C). When all KusakiDB entries were considered the proportion changed to, 91.3% validated by UniProt, 4.5% by RefSeq and 4.2% not validated (Figure 1-D). Among Class and Family taxonomy levels, Liliopsida and Poacea, respectively, were the most represented (Suppl. File 2-A, Suppl. File 2-B).

### Study Case 1: *Triticum aestivum*

Considering the family Poaceae, OG Assessment showed that 50% of *T. aestivum* OGs were in common with 45.45% of other species in the same family (median family), and 9.40% considering all species (median species).

OG management using the criteria KusakiDB ‘Validated’, Total number of species in a family equal to 5, percentage of species in a family and percentage of total species higher than 50%, showed that *T. aestivum* had 4,368 OGs, the lowest number among all families that have at least 5 species in a family (Suppl. File 3-A). A list of not validated OGs using the remaining same parameters can be analyzed on Suppl. File 3-B.

### Study Case 2: Search for potential high evolutionary rate genes

To search for orthologs that are conserved at a family level but not conserved within all species and have a physical evidence of existence (KusakiDB validated), the parameters could be, percentage of total species less than 40% and percentage of species in a family higher than 50%, with total number of species in a family equal or higher than five species. The final result can be downloaded for further analysis (Suppl. File 4).

### User’s Input Data

KusakiDB provides two types of analysis using the output of Hayai-Annotation Plants (Ghelfi, et al., 2019). The details are described in https://github.com/aghelfi/kusakiDB.

## Supporting information

Supplementary_Data.zip

## Acknowledgements

This work was supported by the Life Science Database Integration Project (Database Integration Coordination Program) of the National Bioscience Database Center and Kazusa DNA Research Institute Foundation, Japan. We are grateful to for the fruitful discussions with K. Shirasawa and technical support by H. Hirakawa, M. Yamada and M. Kohara.

## Author Contributions Statement

A. G. conceived and Y. N. and S. I. coordinated the project. A. G., Y. N. and S. I. analyzed and interpreted the data. A. G. developed the software packages. A. G. wrote the manuscript with contributions from Y. N., and S. I. All authors reviewed the manuscript.

